# Sensory adaptation modulates coding and perceptual quality of odor mixtures

**DOI:** 10.1101/2023.09.30.555579

**Authors:** Nicolás Pírez, Federico Andrés Gascue, Fernando Federico Locatelli

**Author notes:** **Corresponding Author:** Dr. Fernando F. Locatelli. Contributed equally (Nicolás Pírez, Federico Andrés Gascue).

## Abstract

The sensitivity of the sensory systems must be dynamic in order to allow animals to adjust their behavior based on experience to optimize detection of relevant information while ignoring stimuli with no predictive value. In this context, one of the main phenomena that modulate the olfactory system is sensory adaptation. It is usually defined as a decrease in the sensitivity or response to a stimulus after a sustained exposure to it. Adaptation may occur in brief intervals of time and depends on the immediate prior experience. Here, we investigate aspects of the function and neurobiology of sensory adaptation in olfaction using the honeybee *Apis mellifera*. By means of electroantennograms we set stimulation protocols that induced sensory adaptation. We show that activation patterns that encode mixtures of odorants in the antennal lobe are drastically altered after sensory adaptation, favoring the representation of stimuli that are present at sub-threshold concentrations. We investigate the effects that sensory adaptation has on the perception of odorant mixtures and show that adapting animals to one of the components of a binary mixture, reduces the appetitive learning of the adapted stimulus and enhances the detection and learning of the non-adapted stimulus in cases in which it would stay normally occluded. These results suggest that olfactory sensory adaptation is critical to allow detection of minor components present in complex mixtures, emphasizing its role as a fundamental mechanism to improve sensitivity to discrete stimuli.

## Introduction

Animals are capable of detecting, processing and learning environmental stimuli, allowing them to cope with changes in their environment. Different sensory modalities play a crucial role in the acquisition of this information, and the mechanisms and computations involved in these processes have been the subject of numerous studies (Chen et al., 2015; Galizia et al., 2000; Locatelli et al., 2016; Locatelli et al., 2013; Sachse and Galizia, 2003). Additionally, animals must be capable of discriminating relevant stimuli immersed in a sea of different stimuli present in the background (Linster et al., 2007; Locatelli et al., 2013). The olfactory system provides us with multiple examples of this, making it an excellent model system to study this question. It is a highly conserved system across different species and most of the neuroanatomical and physiological principles that are involved in the detection, processing and coding of odorant stimuli are very similar between insects and vertebrates (Wilson, 2013; Wilson and Mainen, 2006). Thus, the insect olfactory system has been widely used to study the fundamentals of odor coding. Briefly, odorants are detected by olfactory receptor neurons (ORNs) that send their projections to the antennal lobe (AL), where they make synaptic contacts with second order neurons, projection neurons (PNs) and take part of a complex circuit that also includes local interneurons (LNs). ORNs that express the same receptor protein converge in the AL to form a neuropile called glomerulus (Wilson, 2013; Wilson and Mainen, 2006). Thus, each glomerulus represents a functional unit, of which there are around 160 in the honeybee AL (Arnold et al., 1985; Flanagan and Mercer, 1989) and it has been suggested that their spatio-temporal pattern of activity determines the odorant identity (Kay and Stopfer, 2006; Wilson, 2013; Wilson and Mainen, 2006).

In nature, animals are exposed to complex stimuli that are composed of multiple odorant components, and from this they must be able to detect relevant components that might be occluded by other components of the mixture (Dudareva et al., 2004) (Smith et al., 2006). There are several hypotheses regarding how mixtures of odorants are perceived and processed. In honey bees, this processing is best explained by the unique cue hypothesis where the animals also come up with a supplementary representation for the mixture (Rescorla, 1972; Rescorla, 1973; Whitlow and Wagner, 1972). Here we studied how sensory adaptation might contribute to detect the presence of minor but relevant odors embedded in mixtures. Sensory adaptation is defined as a fast and temporal decrease in the response or sensitivity to a stimulus after a sustained exposure to it (Rankin et al., 2009; Stortkuhl et al., 1999; Twick et al., 2014; Wilson and Linster, 2008). It provides the system with a substrate for plasticity, and it is believed to help in the segregation of stimuli that are present concomitantly, such as mixtures. Some of the characteristics of sensory adaptation are known, such as its rapid recovery after stimulus offset (Cafaro, 2016). It has been suggested to be very important for odor-background segregation in vertebrates (Kadohisa and Wilson, 2006; Linster et al., 2007). In flies, it has been shown that ORNs and PNs adapt when they are stimulated with a constant background odorant (Bhandawat et al., 2007; Cafaro, 2016; de Bruyne et al., 2001; Nagel and Wilson, 2011). However, recordings from these cells have shown that when animals are presented with a novel stimulus that arrives after the onset of the background stimulation, these neurons remain responsive (Cafaro, 2016). In honeybees, the responses of PNs adapt after a short period of odorant presentation (Sachse and Galizia, 2002), suggesting that sensory adaptation could help them discriminate novel odorants on top of tonic odorant backgrounds. It is important to point out that sensory adaptation has been mostly studied focused on the reduction of the response or sensitivity of an animal to a stimulus, but not on the advantages that it can provide to animals, allowing them to detect or perceive stimuli that have not been adapted to. Here we studied the enhancing effect that sensory adaptation has on the ability of animals to detect odor components that normally would be overshadowed by another component. We show that a stimulation of 30 seconds is sufficient to trigger sensory adaptation in honeybees and determine that the activity patterns at the level of the AL are altered after adaptation, making the representation of the mixture more similar to the non-adapted component. Finally, we show that sensory adaptation reduces learning of adapted component, while concomitantly favors learning of non-adapted components that otherwise would stay occluded in the mixture. In summary, we show that sensory adaptation serves to increase the sensibility of the animals to certain stimuli, and not only to decrease it, as normally interpreted.

## Materials and Methods

### Animals

Honey bees (*Apis mellifera*) pollen foragers (all females) were collected, between 10 and 11:30 AM, at the entrance of regular hives located at the campus of the University of Buenos Aires (34° 32 S; 58° 6 W). Once in the laboratory, the bees were briefly cooled and restrained in individual holders (Galizia and Kimmerle, 2004), keeping their proboscis, antennae, and mandibles free. The holders prevented head movements, while leaving the antennae and proboscis free to move. After recovery from cooling, bees were fed 5 µl of a 1 M sucrose solution and remained undisturbed until feeding *ad libitum* in the evening. In the laboratory, the bees were kept in a humid box at room temperature (20°C – 24°C) on a 12:12 h light:dark cycle.

### Electroantennogram

To perform these experiments the body was severed from the head and the antennae were fixed toward the front with Eicosane (Sigma-Aldrich). A silver wire was inserted into the compound eye for ground reference. The most distal segment of the flagellum was cut using a micro-scissor. A glass microelectrode filled with Ringer’s solution (in mM: NaCl, 130; KCl, 6; MgCl_2_, 4; CaCl_2_, 5; sucrose, 160; glucose, 25; and HEPES, 10; pH 6.7, 500 mOsmol) was inserted through the hole at the tip of the antenna, by means of a micromanipulator. Electrical signals passed through a differential amplifier (DAM50, WPI), amplified 10000 times and recorded with a National Instruments data acquisition board (NI USB-6000) connected to a computer for recording and offline analysis. The data was acquired at a 1000 samples/s rate. Recordings were made in one antennae only. In the offline analysis data was filtered with a Gaussian lowpass filter using Clampfit software in order to reduce noise in the signal. There were two types of trials: 1) “control” trials that started with a 31 second baseline, followed by a 4 seconds odorant pulse, lasting 35 seconds; and 2) “adaptation” trials that lasted 60 seconds and were comprised of a 5 seconds baseline, followed by a 30 seconds odorant pulse, a 5 second recovery time, and a final 4 seconds odorant stimulation. The long and short pulses in the adaptation trials could be the same or different odorants depending on the experiment (see Figure 1A and Table 1). All recordings were separated by no less than 1 minute. Odorants used were 1-hexanol and acetophenone (both at 8 % v/v in liquid phase). To quantify sensory adaptation we calculated the amplitude of the response elicited by the 4 seconds odorant pulse in an “adaptation trial”, divided by the amplitude of the response elicited by the 4 seconds pulse of the same odorant in a “control” trial. For amplitude of the response we used the area under the curve.

**Figure 1.**
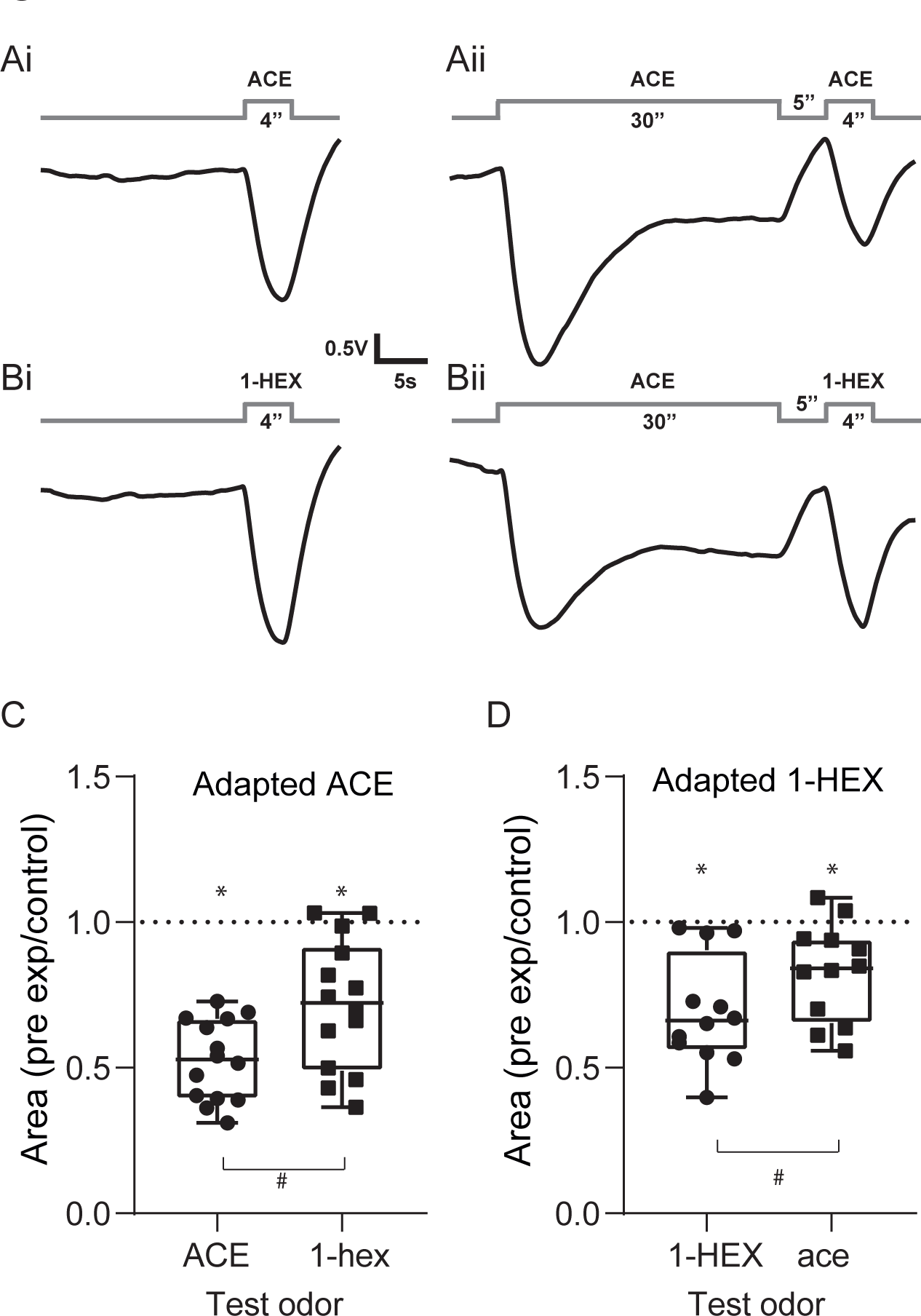
Olfactory adaptation after 30 seconds of exposure to an odorant. *A-B.* Schematic of the stimulation protocol during electroantennogram recordings. **A** and **B** correspond to acetophenone or 1-hexanol used in a counterbalanced way. Left and right panel describe two sorts of trials. ***Ai-Bi*.** Control trials included only a 4-second stimulation with odor and were separated from any other stimulation by not less than 1 minute. ***Aii-Bii***. Adaptation trials consisted of a 30-second odorant stimulation (to trigger adaptation) and a second odorant stimulation (4 sec) separated by 5 sec. In the adaptation trials, the same or a different odorant (as indicated in each case) was used for the pre-exposure and the short stimulation. ***C*.** Sensory adaptation induced by pre-exposure to acetophenone (ACE). Box plots show the sensory adaptation index calculated as the ratio between the amplitude of the odor response during the 4 sec stimulation of an adaptation trial divided by the amplitude of the odor response during the 4 sec stimulation of a control trial. The effect of pre-exposure to acetophenone was measured on acetophenone itself and on 1-hexanol (cross-adaptation). Each condition was first statistically contrasted against the theoretical value of 1 that corresponds to no sensory adaptation (one sample *t-*test, * for p < 0.001). Paired *t-test* was used to compare the effect of pre-exposure to acetophenone on acetophenone itself vs on 1-hexanol (t = 2.667, p = 0.0194, #, N = 14). ***D.*** Same as in **C** but in this case sensory adaptation was induced by exposure to 1-hexanol (1-HEX). One sample *t-*test, * for p < 0.001, for each condition against 1, and paired *t-test* to compare the effect of the pre-exposure on 1-hexanol vs acetophenone (paired *t-test*, t = 2.588, p = 0.0252, #, N = 12).

**Table 1.** Odorant’s concentrations and mixture preparation. Throughout these experiments we used different pure odorant concentrations and different binary mixtures. In the cases in which a binary mixture was used (optical imaging and behavior), one odorant (A) was always in a higher proportion in the mixture than the other component (odorant B).

|  | **Odorant A** | **Odorant B** | **Experimental setup** |
| --- | --- | --- | --- |
| PRE ACE | Acetophenone 8 % v/v | 1-hexanol 8 % v/v | Electroantennogram |
| PRE 1-HEX | 1-hexanol 8 % v/v | Acetophenone 8 % v/v | Electroantennogram |
|  | **Odorant A** | **Odorant B (occluded)** |  |
| PRE ACE 4:1 | Acetophenone 8 % v/v | 1-hexanol 2 % v/v | Optical imaging/Behavior |
| PRE ACE 40:1 | Acetophenone 8 % v/v | 1-hexanol 0.2 % v/v | Optical imaging/Behavior |
| PRE 1-HEX 4:1 | 1-hexanol 8 % v/v | Acetophenone 2 % v/v | Optical imaging/Behavior |
| PRE 1-HEX 40:1 | 1-hexanol 8 % v/v | Acetophenone 0.2 % v/v | Optical imaging/Behavior |
| PRE 1-HEX 200:1 | 1-hexanol 8 % v/v | Acetophenone 0.04 % v/v | Behavior |
| PRE OCT 1:1 | 2-octanone 2 % v/v | Geraniol 2% v/v | Optical imaging/Behavior |
| PRE GER 1000:1 | Geraniol 20 % v/v | 2-octanone 0.02 % v/v | Behavior |

### Projection neuron staining and calcium imaging experiments

Olfactory projection neurons were stained by backfilling with the calcium sensor dye Fura-dextran (potassium salt, 10,000 MW; Invitrogen). Briefly, after capture, the animals were placed in individual holders designed to fit under the microscope. After recovering from chill anesthesia, animals were fed with 4 µl of a 1 M sucrose solution and their head fixed in place with a small drop of wax. A window was cut in the head capsule dorsal to the joints of the antennae and ventral to the medial ocellus. The glands were carefully moved aside until the mushroom body (MB) alpha lobes and calyces were visible. The tip of a glass microelectrode was coated with dextran-conjugated dye and inserted into both sides of the protocerebrum, where the lateral antenna-protocerebral tracts enter the lateral calyces of the MBs. The dye bolus dissolved into the tissue in 3 to 5 seconds. The window was immediately closed using the same piece of cuticle that was previously removed and sealed with low temperature melting Eicosane (Sigma). After staining, the bees were fed with 1 M sucrose solution and left undisturbed until the next day. Before imaging, the antennae were fixed pointing toward the front using Eicosane. The head capsule was opened, and the brain was rinsed with Ringer’s solution. Glands and trachea covering the ALs were removed to facilitate the optical pathway. Only one AL per animal was used for imaging, selecting the one that had more homogeneous staining, by looking at the resting light image (RLI). The bee was placed in the microscope and left undisturbed 15 minutes before the imaging started. Calcium imaging was done using an EMCCD iXon camera (ANDOR) mounted on an upright fluorescence microscope (Olympus BX-50WI) equipped with a 20X dip objective, 0.5 NA (Olympus). Filter and mirror set: 505 DRLPXR dichroic mirror and 515 LP filter (Till Photonics).

Excitation light was provided by a Polychrome V (Till Photonics), which alternated between 340 and 380 nm. Sampling rate was 8 Hz (a frame every 125 msec). Calcium imaging analyses were done using custom software written in Interactive Data Language (IDL; Research Systems) by Giovanni Galizia (University of Konstanz, Konstanz, Germany). Each data set or measurement consisted in a double sequence of 96 images of the antennal lobe, obtained by alternating 340 and 380 nm excitation light (*Fi*_340_, *Fi*_380_, where *i* is the number of the image from 0 to 96). For each pair of images *Fi*, we calculated the ratio *Ri* = (*Fi*_340_ nm / *Fi*_380_ nm) x 100. The resulting values represent the change in fluorescence and are proportional to the changes in the intracellular calcium levels. The analysis of the odor-induced activation patterns in this study was based on the signals from 10 glomeruli (also called ROIs) that were identified on the basis of their morphology and position using published atlas of the honey bee AL (Flanagan and Mercer, 1989; Galizia et al., 1999). The visualization of different ROIs is possible by observing the RLIs obtained at 380 nm excitation light. The glomerular activation was calculated by averaging activity in a square of 7 x 7 pixels that correspond to 23 x 23 μm and fit within the limits of the glomeruli. For some analysis, the temporal detail of the activity was collapsed averaging the response from 1 second (8 frames), starting 2 frames after odorant stimulation onset, which was 4 seconds long and was presented after a baseline period of 3 seconds.

### Odorant Stimulation

For electroantennogram recordings, calcium imaging and behavioural experiments we used the same odorant delivery device. The device consisted of independent channels, each of them attached to a 10 ml glass vial that contained one odorant. The odorants used were 1-hexanol, acetophenone, 2-octanone and geraniol or binary mixtures of them. Inside the vials the odorants were diluted in mineral oil at concentrations that varied depending on the experiment. The concentrations were (v/v): acetophenone 8%, 2%, 0.2% and 0.02%; 1-hexanol 8%, 2% and 0.2%, geraniol 8%; 2-octanone 8% and 0.008%. An additional “empty” channel/vial containing only mineral oil was used as a blank stimulus and to compensate for the flow delivery system in the trials in which only one odorant was being presented. The saturated headspace inside the vials was used as the odor source. The odorant delivery system had a continuous central air stream of 60 ml/s in which the headspace of the vials was injected by nitrogen at a flow of 6 ml/s (N_2_ avoids odorant oxidation). Opening and closing of the odorant channels were controlled by solenoid valves (LFAA1200118H; The LEE Company) and synchronized with the recordings. The odorant concentration that reached the bee resulted in 1/10 of the initial concentration in the headspace of the vial. For stimulation with a pure odorant, the corresponding channel was opened together with the empty channel. For stimulation with a binary mixture, the two corresponding channels were opened, so that the mixture was obtained by mixing the respective headspaces. The longer stimulations used to produce sensory adaptation, were generated by additional channels, to ensure that the main odorant channel for the short stimulations was not depleted (see Supplemental material for controls on this issue). During periods without odorant stimulation the charcoal-filtered air stream continuously ventilated the animal. A gentle exhaust located 10 cm behind the bee removed the odorants from the experimental arena.

### Appetitive olfactory conditioning

All experiments began one day after capture and were performed between noon and 4:00 PM. The day of the experiment, the bees were food deprived a condition that improves appetitive learning. Honey bees were trained using appetitive olfactory conditioning of the proboscis extension reflex (Bitterman et al., 1983; Hammer and Menzel, 1995). During training the conditioned stimulus (CS) was a binary odor mixture and the unconditioned stimulus (US) were 0.2 µl of 2 M sucrose solution applied to the antennae and proboscis. The CS lasted 4 seconds. The US was applied three seconds after the onset of the CS. All conditioning protocols comprised three rewarded trials separated by 10 minutes. The experiments included two different groups of bees. In the “adapted” group, each training trial was preceded by 30 seconds exposure to a pure odorant and 5 seconds resting period before the CS started. The “control” group of bees was trained using the same CS and US, but lacked the exposure that induced adaptation before each training trial. One hour after training, animals were tested by measuring the response to the components of the mixture in separate trials. The concentration of the single odorants during the test was the same that they had in the mixture during the training (see Figure 6A and Table 1). The concentration of the single odorants and the binary mixtures varied across experiments and is indicated in each case.

### Statistical analysis

For the electroantennogram data we performed paired *t* tests to compare the ratio of the response between odorants. Additionally, we analyzed for each odorant if the ratio was different from an expected value of 1 which would correspond to no adaptation. This was done using the one sample *t* test to compare to a theoretical mean of 1. Finally, we calculated a paired pulse ratio to compare the response elicited by the two different 4 second odorant stimulations (A vs. A and A vs. B). This data was also analyzed by a one sample *t* test. For the calcium imaging experiments we performed different analyses. We calculated Pearson correlation coefficients of the odorant evoked patterns of activity, and performed PCA analysis of the responses. When appropriate we used 2-way RM ANOVA, with uncorrected Fisher’s LSD, Sidak’s or Tukey’s post hoc comparisons to compare the different experimental conditions. To test for cross adaptation (Figure 3) among our tested odorants we performed the following analysis: first, we ranked the glomeruli by subtracting their response to odorant B from the response to odorant A. We then proceeded to rank the glomeruli from 1 to 10, with glomerulus 1 being the glomeruli with the highest response to odorant A (either 1-hexanol or acetophenone) and glomerulus 10, the glomeruli with lowest response to odorant B. Next, we normalized the responses of the mix pre and mix adapt before subtracting the first from the second to be plotted as a function of the ranked glomeruli. The rationale behind this analysis was to bring to light the expected effect that highly active glomeruli to an odorant will show a bigger effect (negative values) when they are adapted to that odorant. In the behavioral experiments the individual response of each animal to the different stimuli is a binary response and was recorded as “1” when the animal responded extending its proboscis to the odorant before the reward was presented. We analyzed this data set with mixed effects analysis, since the number of animals was such that allows us to do that, despite being a dichotomic dependent variable. The proportion of positive answers was compared orthogonally between groups using Sidak method (Ludbrook, 1994).

## Results

### Sensory adaptation after 30 seconds of exposure to odor

In this study we evaluated whether and how sensory adaptation to an odorant affects the perceptual quality of a second odorant that is presented shortly after the first one. Thus, our first step was to establish a stimulation protocol able to induce and reveal olfactory sensory adaptation in a reliable and measurable way. For this aim, we measured responses in ORNs by means of electroantennogram recordings. Figure 1 shows the stimulation protocols and EAG examples of a set of experiments performed to measure whether exposure to an odorant reduces the response to an immediate subsequent odorant pulse. Figures 1Ai and Bi correspond to single 4-seconds pulses of acetophenone and 1-hexanol respectively. Figure 1Aii and Bii correspond to EAG recordings that included a 30-seconds odorant stimulation and a 4-seconds stimulation separated by 5 seconds interval. During the 30 seconds stimulation, the EAG recordings peak shortly after odor onset and then their amplitude reaches a plateau a few seconds later, even though the odorant was still present. The recordings show that second pulses elicit smaller antennal responses. To evaluate whether this reduction is specific to the exposed odorant, we performed EAG recordings in which we exposed the bees 30 seconds to acetophenone and measured whether it affected the response to 1-hexanol (Figure 1Bii and D), and vice versa. In order to quantify sensory adaptation, we calculated the ratio between the amplitude of the response elicited by the 4-seconds odor pulse that was preceded by 30 seconds exposure to odorant and the response to a 4-seconds pulse of the same odorant but not preceded by exposure to odorant. As it is summarized in Figure 1C and D, sensory adaptation was more substantial when the odorant used to trigger sensory adaptation was the same that was used for the second pulse. Nonetheless, a certain level of sensory adaptation was also observed when both odorants were different.

### Olfactory sensory adaptation alters the representation of odors in the antennal lobe

Using the sensory adaptation protocol established above, we analyzed whether and how the internal representation of a mixture of odorants is altered when bees are adapted to one of the components of the mixture. In order to measure odor representation at the output of the antennal lobes, we stained uniglomerular PNs by backfilling them with the calcium sensor dye Fura-dextran (Sachse and Galizia, 2003).

All the experiments in this section followed the next design (Figure 2A): first, we measured the activity patterns elicited by stimulation with two pure odorants and their respective binary mixture (mix pre) in trials separated by 60 seconds. Afterwards, the bees were subjected to a 30-seconds exposure to one of the odorants (odor A). Five seconds after offset of the exposure to odor A, the activity elicited by the mixture was measured again (mix adapt). Finally, 60 seconds later, we measured the activity elicited by the mixture for the third and last time (mix recov). Figure 2B shows an example in which the odorant used to induce sensory adaptation was 1-hexanol and the binary mixture was 1-hexanol:acetophenone in proportion 4:1. The false color maps represent calcium imaging patterns elicited by the two odorants and the mixture at the different steps of the experiment. The white squares on the panel to the left show the location of the different regions of interest (i.e. glomeruli) that were analyzed in this example. As observed, the pure odorants activated different but partially overlapping sets of glomeruli. The binary mixture elicited an activity pattern that included all glomeruli activated by the single odorants. After 30 seconds exposure to 1-hexanol, the mixture elicited activity in a combination of glomeruli that resembles the activation pattern elicited by acetophenone alone. After 1 min interval, the initial representation of the mixture was almost completely restored. The activity elicited by the mixture in each glomerulus was in general slightly higher than the activity elicited by the component that elicits the highest activation, indicating a sub-linear summation of the activity elicited by the components (Deisig et al., 2006). The combination of glomeruli activated by the mixture changed drastically after adaptation to 1-hexanol. As observed, the activity pattern elicited by the mixture was drastically reduced, and was basically restricted to the glomeruli that corresponded to acetophenone. After a 60 seconds interval the activity elicited by the mixture was recovered in all regions. Figure 2C shows time resolved heat maps of the same 8 regions of interest from B. Additionally, we performed a principal component analysis (PCA) to represent the complex spatio-temporal patterns of activity elicited by the different odors (Figure 2D, Supplementary Figure 2). The activity patterns elicited upon stimulation with an odorant started from a common origin that corresponds to basal activity before odor onset. Each odorant or mixture describes a distinct trajectory in the PCA space, being 1-hexanol and acetophenone the more dissimilar ones. The activity elicited by the mixture projected within the space delimited by the two pure odorants, but closer to 1-hexanol which in this example was the majority component of the mixture. Immediately after adaptation to 1-hexanol, the trajectory of the activity elicited by the mixture moved drastically towards the direction of the non-adapted odorant, acetophenone. Finally, after a 60 seconds interval, the transient elicited by the mixture recovered a trajectory similar to the one observed before sensory adaptation.

**Figure 2.**
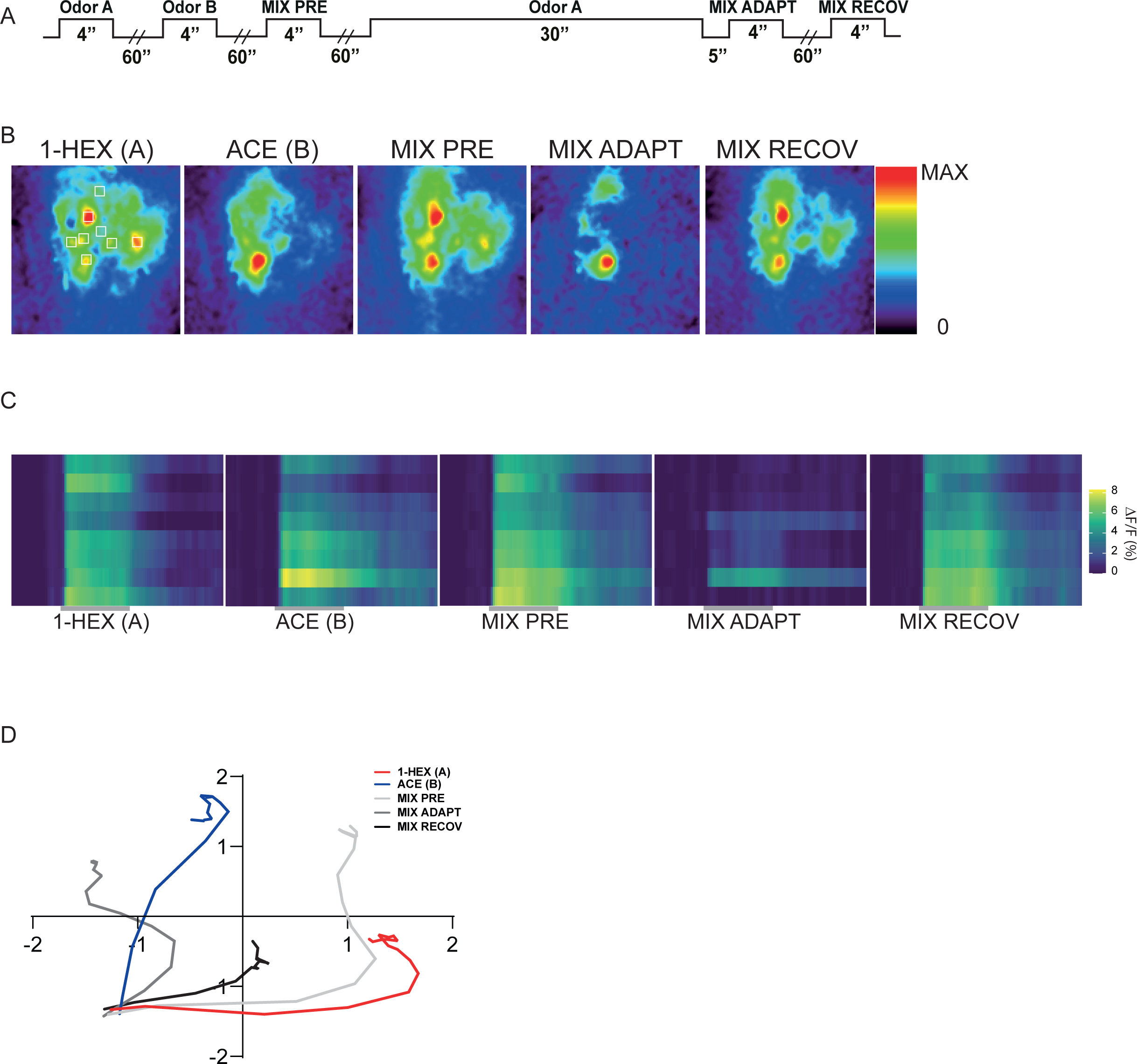
Sensory adaptation alters the representation of mixtures in the antennal lobe. ***A.*** Schematic of the stimulation during optical imaging recordings. A and B correspond to acetophenone or 1-hexanol used in a counterbalanced way. Each imaging trial includes a 4-second stimulation with odor A, followed by a 60 sec interval, then a 4-seconds stimulation with odor B. After another 60 sec interval, the mixture of A and B is presented for the first time (mix pre). Another 60 sec interval follows, and then the animals are exposed for 30 sec to the odor A and 5 sec after this, a second presentation of the mixture (mix adapt). Lastly, following another 60 sec interval, the last stimulation with the mixture (mix recov). ***B.*** Pseudocolor maps showing the peak response in the AL during 4-seconds stimulation measured with calcium imaging. Each picture is autoscaled to show the spatial pattern of activity and not the absolute activity. The white squares show the position of the 8 selected regions of interest (ROIs) on this example. ***C.*** Time resolved heat maps of the same 8 regions of interest from **B**. ***D.*** Principal component analysis showing spatiotemporal patterns elicited bye of the two pure odorants and the mixture (pre, adapt and recov).

### Cross-adaptation from components to mixtures

So far, we have observed that sensory adaptation to an odorant causes transient changes in the representation of a mixture that contains this odorant. In the next section, we aimed at analyzing these changes at the level of glomeruli. For that aim, we selected in each animal 10 ROIs from the ventral side of the antennal lobe that based on their size and activation profiles correspond to 10 glomeruli. Each ROI/glomerulus was ranked according to whether it was activated by the “to be used for adaptation” component (odor A) or by the “control” component of the mixture (odor B). We made this ranking by first normalizing the pattern of activity elicited by each odorant in each bee and then subtracting the normalized activity elicited by odorant B from the activity elicited by odorant A in each glomerulus. This procedure provided an index for each glomerulus, in which higher values correspond to glomeruli that are activated by odorant A but not B, lower values correspond to glomeruli that are activated by odorant B but not A, and intermediate values indicate glomeruli that were activated by both of them (Figure 3). This classification allowed us to averaged glomeruli from different animals on the basis of their response towards the components of the mixture and analyzed their respective impact on the changes observed after sensory adaptation. Figure 3A corresponds to the results of a first group of bees in which odorant A was acetophenone and odorant B was 1-hexanol. The mixture was built in proportion 4:1; acetophenone:1-hexanol, and acetophenone was used as the adapting odorant. Notice that in all experiments, odorant A was always used in higher concentration than odorant B, since the mixtures were designed to make A the dominant component of the mixture. Figure 3A shows for each glomerulus the absolute difference between the activity elicited by the mixture before and after sensory adaptation to odor A. From left to right, the glomeruli are ordered based on the response ranking from odorant A to odorant B. As observed, the changes in the representation of the mixture after adaptation are mainly explained by a reduction in the activity of the glomeruli that are activated by acetophenone (i.e. odor A). Figures 3B, C and D correspond to other three sets of experimental bees in which the role of acetophenone and 1-hexanol as the adapted and control components were alternated, and also different proportions of both odorants were tested. In all cases we observed a similar graded effect in which the glomeruli activated by the adapted odorant are more heavily affected and thus they are responsible for changes in the representation of the mixture. In all cases we calculated the linear best fit for these data sets and found the slope to be significantly different from zero (p value < 0.001, see figure legend for more statistical information) regardless of the odorant used for adaptation and their proportions.

**Figure 3.**
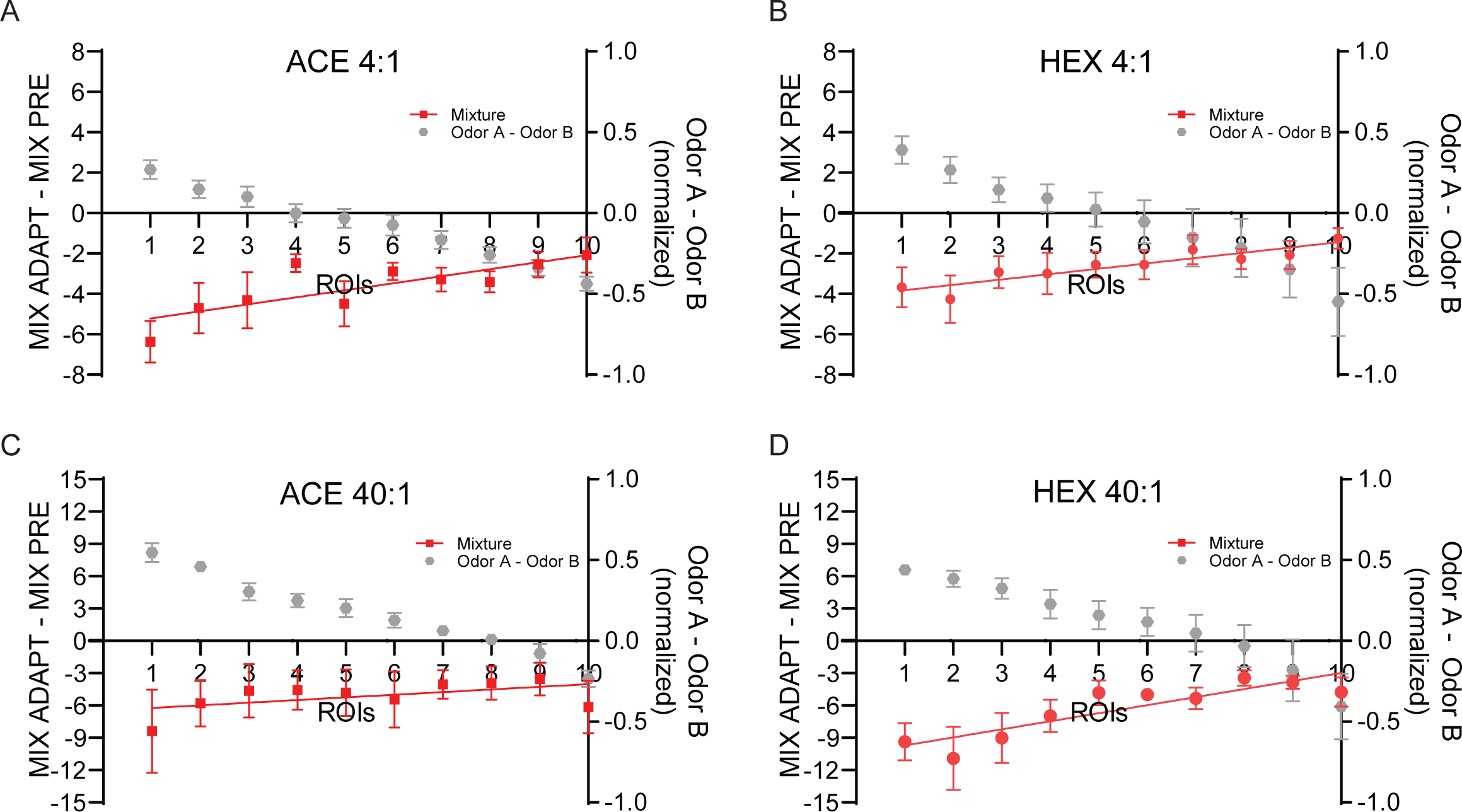
Cross adaptation between components and mixture analyzed at the level of glomeruli. Difference between the activity elicited by the mixture before and after sensory adaptation. Glomeruli are ordered from left to right according to their specificity to odorant A (adapted odor) and odorant B (control odor). Gray circles: average across bees of the index used to categorize glomeruli according to their specificity. Index value is indicated on the Y axis at the right side of each panel. Red circles: Mean and SEM of the change in the response to the mixture (mix adapted minus mix pre). **A.** Bees evaluated with acetophenone and 1-hexanol at ratio 4:1; acetophenone used for sensory adaptation. N = 10. Line: best fit with a significance level of p=0.0007: y=0.3482*X - 5.567, R^2^=0.1388. **B**. 1-hexanol and acetophenone; ratio 4:1; 1-hexanol used for sensory adaptation. N = 10. Line: p=0.0031: y= 0.2654*X - 4.097, R^2^=0.1236. **C**. acetophenone and 1-hexanol; 40:1; acetophenone used for sensory adaptation. N = 10. Line p=0.3084: y=0.2458*X - 6.477, R^2^=0.0216. **D**. 1-hexanol and acetophenone; 40:1; 1-hexanol used for sensory adaptation. N = 10. Line p < 0.0001: y= 0.7479*X - 10.46, R^2^=0.4235.

### The correlation between the patterns elicited by the mixture and the pure components changes during sensory adaptation

To quantify the changes at the level of the neural population response that encodes the mixture before and after adaptation, we calculated the Pearson correlation coefficients between the glomerular activity patterns elicited by the pure components and the mixture before and after adaptation. Figure 4A summarizes the results of the first group of bees in which acetophenone was used as odorant A (adapted), 1-hexanol as odorant B (non-adapted), and the mixture ratio was 4:1 (ace:1-hex). As observed, the correlation between the glomerular activity patterns elicited by acetophenone and the mixture was severely affected 5 seconds after the end of the 30-seconds exposure to acetophenone. After an interval of 1 minute, the initial correlation value was almost completely restored (2-way RM ANOVA: treatment: F_(2, 28)_ = 30.60, p < 0.0001, N = 8, different uppercase letters represent significant differences by Tukey’s multiple comparisons test). The correlation between the non-adapted component, 1-hexanol, and the mixture was also affected, however to a minor extent (2-way RM ANOVA: treatment: F_(2, 28)_ = 30.60, p < 0.0001, N = 8, different lowercase letters represent significant differences by Tukey’s multiple comparisons test). This effect can be explained by the fact that 1-hexanol and acetophenone share part of the glomeruli in their respective activation patterns, as it is evident by the high correlation coefficient between them, also indicated in the figure (blue triangle). Thus, we can infer that exposing the animals to acetophenone induces adaptation also in glomeruli that are shared with 1-hexanol, producing the change in the correlation between the mixture and the non-adapted component too. In a second group of bees, we used 1-hexanol as odorant A and acetophenone as odorant B (Figure 4B). We found that the correlation between the pattern elicited by 1-hexanol and the mixture was drastically affected during adaptation and it was almost completely recovered after 1 minute (Figure 4B, 2-way RM ANOVA, interaction: F_(2,24)_ = 19.07, p < 0.0001, N = 7, different lowercase letters represent significant differences by Tukey’s multiple comparisons test). Interestingly, the adaptation to 1-hexanol did not negatively affect the correlation between the mixture and the non-adapted odor acetophenone. Instead, it showed a slight tendency for increase. This observation constitutes a very interesting phenomenon, since it shows that adaptation to one component of the mixture can potentially enhance the representation of non-adapted components in the mixture. This time, the correlation coefficient between the activity patterns elicited by the two single odorants was lower than in the previous combination, indicating that the overlap between the respective glomerular patterns was smaller, which explains that the correlation between the non-adapted odor and the mixture was less affected. Notice here that the correlation coefficient between 1-hexanol and acetophenone is different between the first and second groups of bees since the ratio acetophenone:1-hexanol was 4:1 in one case and 1:4 in the other.

**Figure 4.**
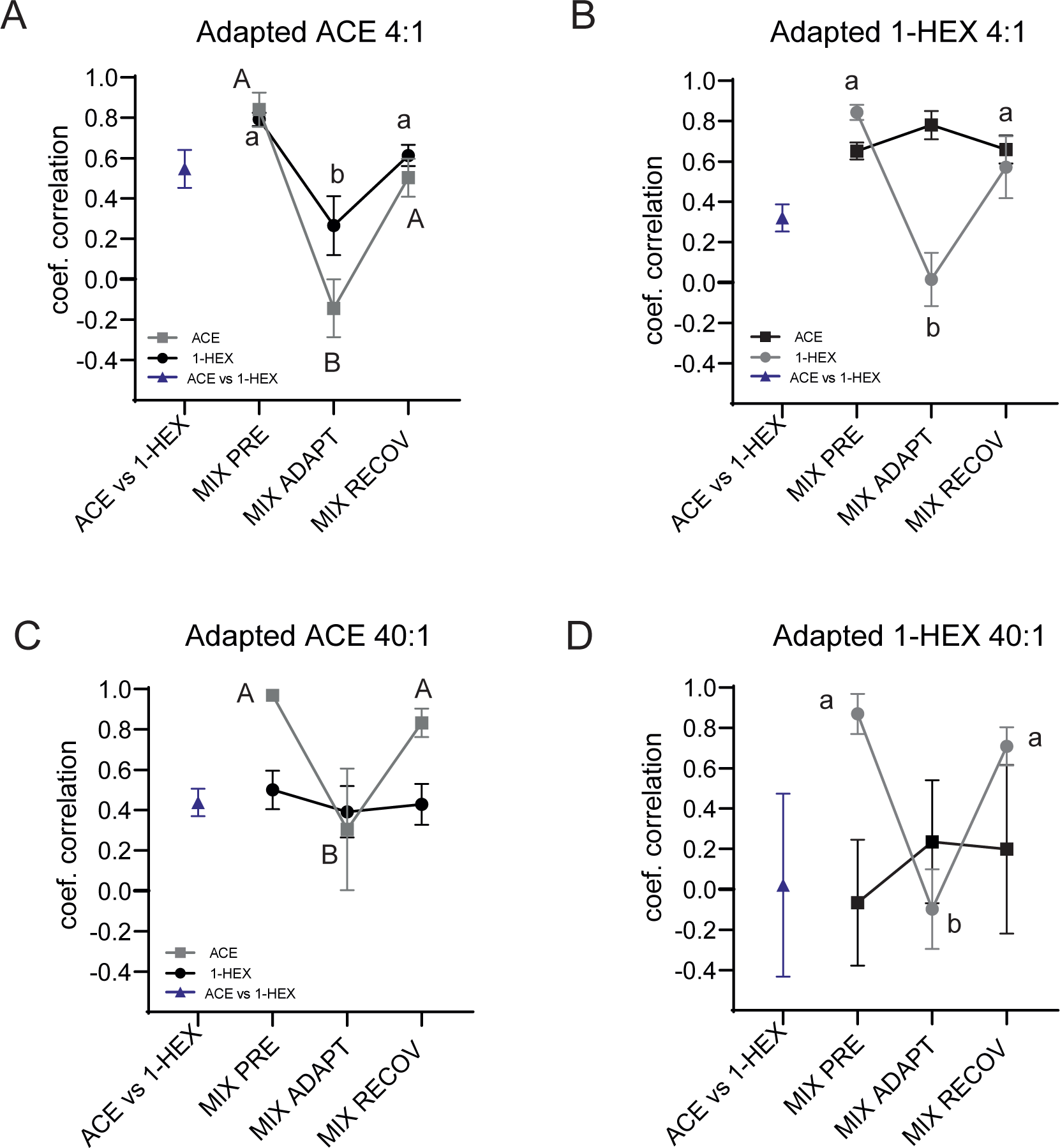
Sensory adaptation modifies the relative contribution of the components to the representation of the mixture. Pearson correlation coefficient between the patterns of activity elicited by the two pure odorants and the mixture, before, during and after recovery from sensory adaptation to one of the odorants. Mean and SEM in all cases. Gray symbols: adapted component. Black symbols: non-adapted component. Blue symbols: Correlation among patterns of activity elicited by the pure components ***A.*** Mixture ace:1-hex, ratio 4:1 and adaptation to acetophenone (2-way RM ANOVA: treatment: F_(2, 28)_ = 30.60, p < = 0.0001, N = 8, different letters represent significant differences by Tukey’s multiple comparisons test). ***B.*** Mixture 1-hex:ace, ratio 4:1, adaptation to 1-hexanol (2-way RM ANOVA, interaction: F_(2,24)_ = 19.07, p < 0.0001, N = 7, different letters represent significant differences by Tukey’s multiple comparisons test). ***C.*** mixture ace:1hex, ratio 40:1, adaptation to acetophenone (2-way RM ANOVA, treatment: F_(2,16)_ = 4.989, p = 0.0207, N = 5, different letters represent significant differences by Tukey’s multiple comparisons test). ***D.*** Mixture 1-hex:ace, ratio 40:1, adaptation to 1-hex (2-way RM ANOVA, interaction: F_(2, 8)_ = 8.842, p = 0.0094, N= 3, different letters represent significant differences by Tukey’s multiple comparisons test).

In the next two experiments we changed the ratio between odorants A and B to 40:1 and 1:40 respectively. Figure 4C shows the data of an experiment in which acetophenone was used as odorant A and 1-hexanol as B. As expected, reducing the relative concentration of the minority component 1-hexanol did also reduce its contribution to the pattern of activity elicited by the mixture, which is observed as a lower correlation value among the patterns elicited by 1-hexanol and the mixture at the beginning of the experiment. Thirty seconds exposure to acetophenone clearly diminished the correlation value between the patterns elicited by acetophenone and the mixture (2-way RM ANOVA, treatment: F_(2,16)_ = 4.989, p = 0.0207, N = 5, different uppercase letters represent significant differences by Tukey’s multiple comparisons test), however did not affect the correlation between the minority component 1-hexanol and the mixture, which was stable across the experiment. Finally, when 1-hexanol was used in higher relative concentration (Figure 4D), we found that 1-hexanol dominated the activity pattern elicited by the mixture (Rhex-mix = 0.9) while the presence of acetophenone was occluded. The reduction of the correlation between the adapted odorant and the mixture was evident as in the previous experiments (2-way RM ANOVA, interaction: F_(2, 8)_ = 8.842, p = 0.0094, N = 3, different lowercase letters represent significant differences by Tukey’s multiple comparisons test). Interestingly, acetophenone showed a tendency to increase its correlation with the mixture following the adaptation to 1-hexanol. One minute later the correlation between both components and the mixture recovered their initial values.

### Sensory adaptation reduces learning of the pre-exposed component of the mixture

Next, we evaluated whether the changes observed in the representation of the mixture correlate with changes in the ability of animals to detect the components that are embedded in the mixture. For this aim, we used appetitive olfactory conditioning of the proboscis extension response (Bitterman et al., 1983). The experimental design consisted of three appetitive conditioning trials using a binary mixture as conditioned stimulus that anticipates a sucrose reward. Each training trial was preceded by 30 seconds adaptation to one of the components of the mixture. One hour after training the conditioned response was evaluated by using the two components in separate test trials. A control group of bees was trained and tested in the same way, but lacked the adaptation to odor before each training trial (see Figure 5A).

**Figure 5.**
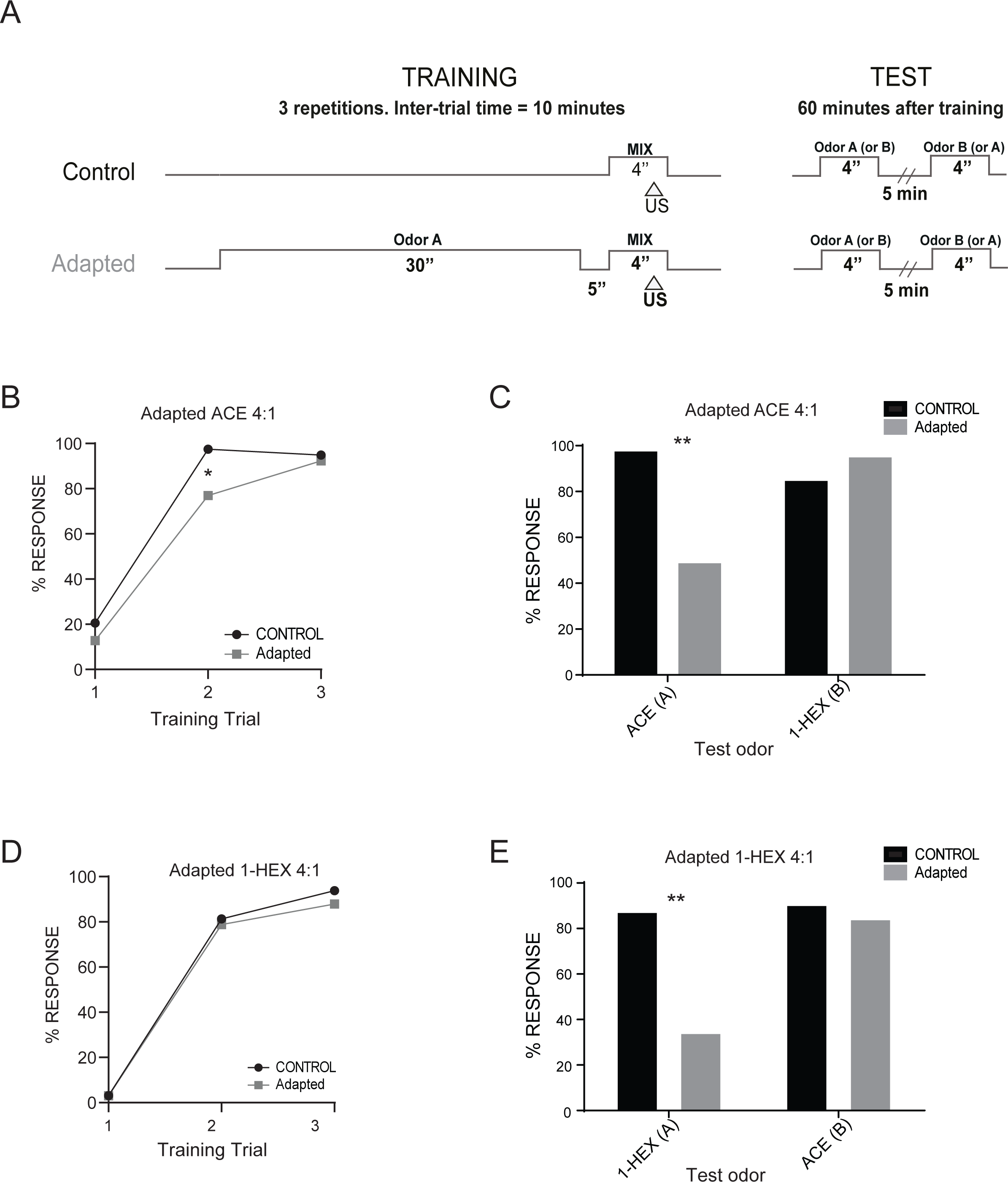
Sensory adaptation to one of the components decreases its learning when the mixture is rewarded. ***A.*** Schematic of the behavioral protocol. Each experiment consisted of an “adapted” and “control” group of bees. Odorants A and B were 1-hexanol or acetophenone as indicated in each experiment. In all cases odorant A is the majority component and was used to trigger sensory adaptation. Odorant B is always the minority component. The mixture of A and B contained as much of odorant A and odorant B as when used alone. US indicate the presentation of the sucrose reward during the CS. Each animal received 3 conditioning trials separated by an ITI of 10 min. In the adapted group, each trial was preceded by 30 seconds exposure to odor A. An hour later, the animals were tested with both components in separate trials separated by 5 minutes, in pseudorandom order and at the same concentration used during training. ***B.*** Training curves of a group of bees in which odorant A was acetophenone 8% and odorant B was 1-hexanol 2%. Animals were adapted to acetophenone. The graph shows the training sessions expressed as the percentage of bees that extend the proboscis upon stimulation with the mixture and before stimulation with sucrose during the training trials. The response of each group to the odorants was analyzed by means of 2-way Mixed Effects Analysis (group: F_(1,76)_=4.661, p=0.034) followed by Sidak’s post hoc test (* represents a significant difference of p < 0.05). ***C.*** The panel shows the percentage of bees that extend their proboscis upon stimulation with the individual components during the testing session (2-way Mixed effects Analysis interaction: F_(1,76)_= 36.68, p < 0.001, N = 39). ***D***. Same as in B but in these experiments 1-hexanol 8% was odorant A and acetophenone 2% was odorant B, thus animals were adapted to 1-hexanol (2-way Mixed effects Analysis, group: F_(1,63)_= 0.256, p=0.61, N = 39). ***E.*** The graph shows the percentage of bees that extend the proboscis upon stimulation with the components during testing (2-way Mixed Effects Analysis interaction: F_(1,62)_=19.87, p<0.001, N = 32). The response of each group to the different odorant was analyzed by means of Sidak’s post hoc test (* represents a significant difference of p<0.01).

Figure 5B corresponds to the first set of bees in which we used a mixture composed of acetophenone and 1-hexanol in a ratio of 4:1 respectively. Acetophenone was used to induce sensory adaptation before each training trial. As it is observed from the performance during training trials, the bees were able to learn the anticipatory value of the mixtures, which indicates that the salience of the mixture as a whole was not affected by adaptation to the component acetophenone. However, the adapted animals reached the higher level of response only at the third trial, having a significantly lower response during the second trial when compared with the control animals (2-way Mixed-Effects Analysis: group: F_(1,76)_= 4.661, p = 0.034; Sidak’s post hoc test of trial 2: p = 0.015). Interestingly, testing the animals with the components separately reveals that sensory adaptation changed the contribution of the components to the perceptual quality of the mixture (Figure 5C). The percentage of bees that showed conditioned response upon testing with acetophenone was significantly lower in the adapted than in the control group (black bars, 2-way Mixed-Effects Analysis: interaction: F_(1,76)_= 36.68, p < 0.001; Sidak’s post hoc test p < 0.01). On the other hand, the response to 1-hexanol was high and did not differ between groups, suggesting that the adaptation was specific to the adapted component of the mixture. Next, we interchanged the roles of the odorants and performed an experiment in which the adapting component was 1-hexanol (Figure 5D). In contrast with the previous experiment, the training curves did not differ between control and adapted groups, and the differences between groups emerged during the test trials with the components. The bees that were adapted during training showed a significant lower conditioned response to 1-hexanol consistent with the interpretation that perception of 1-hexanol was diminished during the training trials (Figure 5E, 2-way Mixed effects Analysis: interaction: F_(1,62)_= 19.87, p < 0.001; Sidak’s post hoc test p < 0.01). On the other hand, no difference among groups was observed for the non-adapted component acetophenone. In conclusion, since the test trials were not preceded by exposure to the odorant and were similar in control and adapted groups, the difference disclosed during the test must have emerged from differences in the way the animals perceive the mixture during the training trials.

### Sensory adaptation and detection of the minor components

In the following experiments we asked whether a minority component of a mixture whose presence would go undetected under normal conditions can be noticed if the animal is subject to sensory adaptation to the dominant component (i.e. the majority component of the mixture). Based on the performance of the control groups in the two previous experiments we concluded that the 4:1 ratio between acetophenone and 1-hexanol was not sufficiently unbalanced to avoid perception and learning of the less concentrated odorant. Thus, in the next series of experiments, we increased the ratio between the components by reducing ten times the concentration of the minor component.

Figure 6A shows the results obtained when acetophenone was used to trigger sensory adaptation and it was 40 times more concentrated than the minority component, 1-hexanol. The training curves (graph on the left) show that even though the control group reaches its highest percentage of conditioned response in the second training trial, the adapted group reached this level in the third trial, indicating the weakening of the salience of the mixture (2-way Mixed-Effects Analysis: group: F_(1,68)_= 8.435, p < 0.01; Sidak’s post hoc test of trial 2: p < 0.01). The subsequent test trials also revealed differences between groups in regard to the perceptual quality of the mixture (bar graph on the right). The control group showed a marked difference in the percentage of conditioned responses evoked by acetophenone and 1-hexanol which reflects the difference in their concentrations. Strikingly, the adapted group shows a significantly higher percentage of conditioned response elicited by 1-hexanol compared with the control group (2-way Mixed Effect Analysis: interaction: F_(1,68)_= 26.31, p < 0.001; Sidak’s post hoc test p < 0.01). This result is important twofold; first because it confirms that 1-hexanol can be detected and learned at this concentration, and second it shows that learning the minority component was facilitated when the animal was adapted to the dominant component during the training trials. Interestingly, this time, sensory adaptation to acetophenone was not evidenced as a lower conditioned response to this odor, probably because its relatively high concentration helps to make it salient enough to be learned. Thus, losing the ability to overshadow a competing odorant might result in a more sensitive characteristic of sensory adaptation than a decreased response to the adapted odor itself.

**Figure 6.**
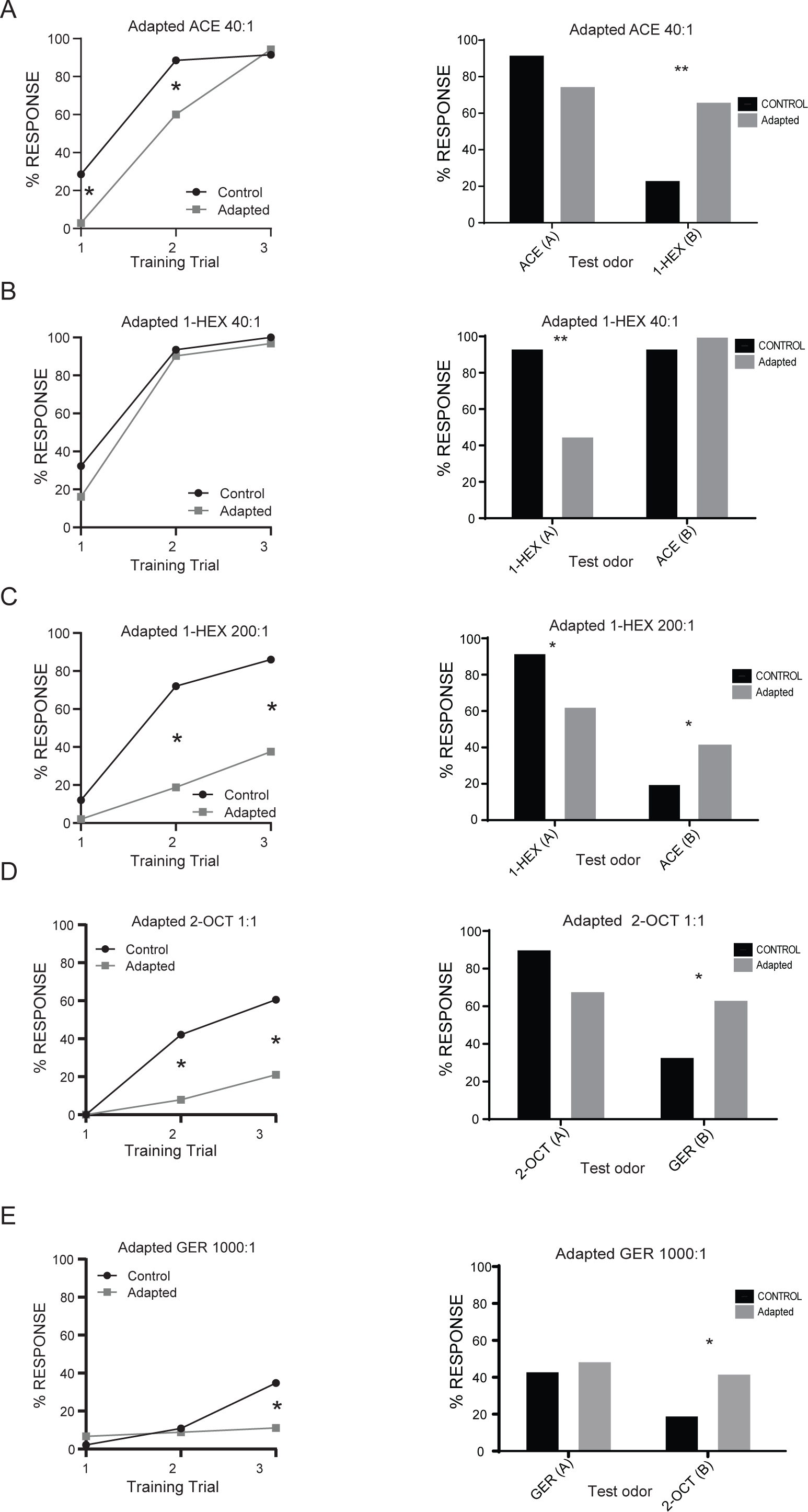
Sensory adaptation unmasks overshadowed components of the mixture. Experimental design and groups as in Figure 5. **Left panels:** Percentage of bees extending the proboscis during training trials upon stimulation with the mixture and before stimulation with sucrose. **Right Panels:** Percentage of bees extending the proboscis during test trials with the components one hour after training. In all cases, black corresponds to control groups and gray to adapted groups. ***A.*** acetophenone 8% and 1-hexanol 0.2%, bees were adapted to acetophenone 8% before each training trial. The response of each group to the odorants was analyzed by means of 2-way Mixed Effects Analysis (interaction: F_(2,136)_=5.869, p < 0.01) followed by Sidak’s post hoc test (* represents a significant difference of p < 0.05). Test session: 2-way Mixed Effects Analysis interaction: F_(1,68)_ = 26.31, p < 0.001, N = 35, ** represents a significant difference, p < 0.01, Sidak’s post hoc test for response of the groups to the indicated odor. ***B.*** 1-hexanol 8% and acetophenone 0.2%, bees adapted to 1-hexanol 8% before each training trial. The response of each group to the odorants was analyzed by means of 2-way Mixed Effects Analysis (group: F_(1,60)_=2.509, p=0.119) followed by Sidak’s post hoc test). Test session: 2-way Mixed Effects Analysis interaction: F_(1,60)_ = 28.90, p < 0.001, N = 31, * represents a significant difference, p < 0.01, Sidak’s post hoc test for response of the groups to the indicated odor. ***C.*** 1-hexanol 8% and acetophenone 0.04%, bees adapted to 1-hexanol 8% before each training trial. The response of each group to the odorants was analyzed by means of 2-way Mixed Effects Analysis (interaction: F_(2,192)_=13.15, p<0.01) followed by Sidak’s post hoc test (* represents a significant difference of p < 0.05). Test session: 2-way Mixed Effects Analysis (interaction: F_(1,96)_ = 30.78, p < 0.001), N = 50 for control group and 48 for adapted group, * represents significant difference, p < 0.05, Sidak’s post hoc test for response of the groups to the indicated odor. ***D.*** 2-octanone 8% and geraniol 8%, bees adapted to 2-octanone before each training trial. The response of each group to the odorants was analyzed by means of 2-way Mixed Effects Analysis (interaction: F_(2,148)_=9.34, p<0.01) followed by Sidak’s post hoc test (* represents a significant difference of p < 0.05). Test session: 2-way Mixed Effects Analysis (interaction: F_(1,74)_ = 29.02, p < 0.001, N = 38, * represents significant differences, p < 0.05, Sidak’s post hoc test for response of the groups to the indicated odor. ***E.*** geraniol 8% and 2-octanone 0.08%, bees adapted to geraniol 8% before each training trial. The response of each group to the odorants was analyzed by means of 2-way Mixed Effects Analysis (interaction: F_(2,178)_=6.956, p<0.01) followed by Sidak’s post hoc test (* represents a significant difference of p < 0.05). Test session: 2-way Mixed Effects Analysis (interaction: F_(1,89)_ = 2.60, p = 0.11), N = 46 for control group and 45 for adapted group, * represents a significant difference of p < 0.05 Sidak’s post hoc test for response of the groups to the indicated odor.

Next we performed a mirror experiment in which 1-hexanol was 40 times more concentrated than acetophenone and it was used for sensory adaptation (Figure 6B). As observed, the training curves did not differ between groups. During testing, the adapted group showed a significant lower conditioned response upon testing with 1-hexanol compared to the control group (2-way Mixed Effects Analysis: interaction: F_(1,60)_= 28.90, p < 0.001; Sidak’s post hoc test p < 0.01) while the conditioned response to the minority component acetophenone did not differ between groups. These results replicate those obtained in the experiments based on the ratio 4:1 and thus indicate that the presence of acetophenone can be detected and learned despite this relatively low concentration in relation 1-hexanol.

Thus, in the next experiment, we increased the ratio between 1-hexanol and acetophenone to 200:1, in order to obtain a condition in which 1-hexanol is able to overshadow the presence of acetophenone (Figure 6C). This time, the training curves were very different between the control and adapted groups. While the control group showed an acquisition curve similar to those of control groups in previous experiments, the acquisition was drastically affected in the adapted group. This reduction can be explained by sensory adaptation to the majority component together with the lower concentration of acetophenone which this time was not able to compensate for the weakening of the mixture (2-way Mixed-Effects Analysis: interaction: F_(2,192)_ = 13.15, p < 0.01; Sidak’s post hoc test of trials 2 and 3: p < 0.01). The subsequent test trials additionally revealed that sensory adaptation altered the weight of the components in the perception of the mixture. As observed in figure 6C, the conditioned response elicited by the minority component acetophenone was higher in the adapted group than in the control group (2-way Mixed Effects Analysis: interaction: F_(1,96)_= 30.78, p < 0.001; Sidak’s post hoc test p = 0.0245). Thus, sensory adaptation to the major component 1-hexanol rescued the learning of the minority component acetophenone.

Finally, we extended the experiments to a new pair of odorants, 2-octanone and geraniol. Since we knew from preliminary investigations that 2-octanone is learned faster and elicits stronger responses in the antennal lobe than geraniol, we started mixing them at the same concentration of 8% and considered 2-octanone as the dominant component to test the effect of sensory adaptation. As shown in Figure 6D, the training curve was severely affected in the adapted group, which is consistent with a reduced salience of the mixture during training (2-way Mixed-Effects Analysis: interaction: F_(2,148)_= 9.34, p < 0.01; Sidak’s post hoc test of trials 2 and 3: p < 0.01). The subsequent test trials showed that sensory adaptation had opposite effects on learning of each component. While it reduced the learning of 2-octanone, it facilitated the learning of geraniol (2-way Mixed Effects Analysis: interaction: F_(1,74)_= 29.02, p < 0.001; Sidak’s post hoc test p = 0.0244). These results support the notion that the reduced learning to geraniol in the control group cannot be explained by a lower salience of this odor alone, but rather by a certain level of overshadowing caused by 2-octanone. In turn, when the weight of 2-octanone is reduced by sensory adaptation, the perception of the minor component geraniol is enhanced and its learning is facilitated. To evaluate the same two odors but in inverse roles, we used a mixture of geraniol 8% and 2-octanone 0.008%. These proportions were set to compensate for the difference in salience and to boost geraniol to overshadow 2-octanone in control conditions. The slow learning curve of the control group is consistent with the low salience of geraniol and the low concentration of 2-octanone. Indeed, the learning was not evident when animals were adapted to geraniol (2-way Mixed-Effects Analysis: interaction: F_(2,178)_ = 6.956, p < 0.01; Sidak’s post hoc test of trial 3: p < 0.01). Nonetheless, the positive effect that sensory adaptation has on the learning of the minor component was clearly observed during the testing (2-way Mixed Effects Analysis: interaction: F_(1,89)_= 2.60, p = 0.11; Sidak’s post hoc test p = 0.0497). In summary, our results show that the effect of sensory adaptation can be seen both, at the level of behavior and at the neural representation of olfactory stimuli in the antennal lobe. Following a stimulus that triggers adaptation, the perception of odors is altered in a way that the detection of the adapted components is reduced while overshadowed components are unmasked. The phenomenon of unmasking enhances the capability of animals to detect odorants that even in very low concentrations can be detected if they are presented in a timely discrete manner.

## Discussion

Sensory adaptation is canonically defined as the phenomenon by which the sensitivity to a stimulus decreases during persistent exposure to it and is fully recovered in a short period of time after stimulus offset. The consequence that sensory adaptation to a given stimulus has on the perception of other stimuli for which the animal is not adapted has not been thoroughly addressed. Here we analyze and show that sensory adaptation to an odorant facilitates the perception of other non-adapted odorants that are presented concomitantly with the first one and that would stay occluded if sensory adaptation did not mediate.

### Sensory adaptation dynamics

Previous studies have shown in flies *Drosophila melanogaster* that odor pulses of 15 seconds or longer elicit olfactory sensory adaptation that is observed both, at the level of the olfactory sensory neurons responses (Cafaro, 2016) and at the level of odor elicited behavior (Stortkuhl et al., 1999). Here, we performed electroantennogram recordings, and confirmed that an odorant stimulation of 30 seconds was sufficient to trigger olfactory adaptation in honey bees *Apis mellifera*. A more exhaustive search based on different exposures times, concentrations and recovery times will provide a more detailed description of sensory adaptation dynamics in honey bees, nevertheless this result served us as a solid platform for the rest of the imaging and behavior experiments. The antennal electrical response elicited by a 4-seconds odor pulse, was reduced to approximately 50% when it was preceded by 30 seconds of stimulation with the same odor and a 5-seconds recovery interval. In addition, we found a smaller but significant amount of cross-adaptation among the two odorants we tested: 1-hexanol and acetophenone. The calcium imaging recordings, which allowed visualizing the spatiotemporal patterns of activity elicited in the antennal lobe by each odorant, showed a partial overlap among the corresponding activation patterns that is consistent with a certain level of cross adaptation at the level of sensory neurons.

### Odors representation in the AL is altered by sensory adaptation

Relevant olfactory cues that trigger and guide behaviors in natural conditions come normally mixed with background odors that can reduce or even occlude the detection of the relevant ones(Conchou et al., 2019; Dupuy et al., 2017; Rusch et al., 2016). For example, a specific floral odor that may signal a profitable nectar source for honey bees can easily get mixed in turbulent plumes with volatiles emanating from non-informative parts of the plant or from nearby plants of other species. In this context, being able to learn and detect the components of a mixture that is a reliable food predictor, constitutes a challenging task that involves sensory processing, adaptation and plasticity at more than one level along the olfactory circuit (Marachlian et al., 2021; Root et al., 2007; Sehdev and Szyszka, 2019; Smith, 2012). Our main motivation in the present study was to evaluate how sensory adaptation contributes to modulate the perception of complex stimuli in a way that it can enhance detection of relevant cues that are present at minor (underrepresented) concentrations. For that aim we based our experiments on asymmetric binary mixtures in which one of the components was present in a low concentration that makes it difficult to be detected when mixed with a second component at higher concentration. By using calcium imaging of odorant evoked activity in the antennal lobe, we were able to identify the activation patterns elicited by each individual component and how they contribute to the representation of the mixture (Chen et al., 2015; Deisig et al., 2010). Our results show that the glomerular response patterns encoding the mixture are drastically altered by sensory adaptation to one of the components, in a way that it reduces the contribution of the adapted component and enhances the contribution of the non-adapted component. The changes in the representation of the mixture were brief, since 60 seconds after the end of the stimulation that evoked sensory adaptation the recovery of the representation of mixtures was almost complete. Such a fast recovery is consistent with a fast and transient effect of sensory adaptation, rather than with stable changes that are elicited when animals are taught to actively ignore an odor that is repeatedly presented without any contingency (Locatelli et al., 2013). In this latter case, the mixture representation moved in the same direction as shown here, however the changes lasted for at least 70 min after learning and arised entirely in the antennal lobe as the changes were not accompanied by decreased activity in sensory neurons (Locatelli et al., 2013).

We found that adaptation to acetophenone and 1-hexanol did not produce exactly symmetrical effects. Generalization was clearer from acetophenone to 1-hexanol than in the opposite direction, both in EAG recordings and in the calcium imaging experiments. In addition we found that acetophenone was more resilient to occlusion by 1-hexanol than the other way around (e.g. 1-hexanol by acetophenone). These asymmetries might represent differences in odor salience and asymmetric overlap among their respective population codes at the receptor and the central brain levels. Despite these differences, it is important to highlight that sensory adaptation worked in similar ways independently of the odor. These experiments suggest that sensory adaptation modulates the response elicited by sensory stimuli at the central nervous system, in addition to the periphery.

### Odorant input modulates sensory adaptation

Additionally we found that the input to the AL has the potential to modulate the degree of sensory adaptation suffered by the different ORNs. One hypothesized role of presynaptic inhibition at the synapse between ORNs and PNs is to function as a gain control mechanism in which strong odorant-evoked inputs are preferentially suppressed (McGann et al., 2005; Nickell et al., 1994; Olsen and Wilson, 2008; Pírez and Wachowiak, 2008; Wachowiak and Shipley, 2006). We wonder if sensory adaptation might be performing a similar role, where the input to the different glomeruli could determine the degree of adaptation suffered. To test this hypothesis, we used the difference in the response to the mixture after adaptation relative to the response before adaptation as a measure of the strength of adaptation suffered by each glomeruli. After ranking the glomerular responses as a function of the response to the individual odorants, we found that adaptation was stronger on glomeruli that responded more to the odorant used to trigger adaptation, thus strongly activated glomeruli undergo a stronger reduction of its response. This suggests that sensory adaptation could act as a filter to facilitate the detection of odorants that activate glomeruli in a weakly fashion, especially in conditions in which their response would go unnoticed.

Moreover, we can conclude from our data that the phenomenon of olfactory sensory adaptation plays a larger than anticipated role in the detection of odorants present in binary mixtures. The effect can be seen both in the electrical responses at the level of the ORNs, as well as at the level of the first synapse on the olfactory pathway. The neural representation of the stimuli following adaptation can be modulated, and this phenomenon can be both generalized and have a cross-over effect on the response to different odorants. The representation of odorant mixtures is altered in a way that is not only reducing the detection of adapted odor, such as an irrelevant background stimulus, but is also unmasking overshadowed components and by that is enhancing the capability of the animals to detect relevant stimuli.

### Olfactory sensory adaptation modulates the behavioral response

To be able to show that a phenomenon like sensory adaptation has a role in the detection and perception of odorants that initially do not evoke a behavioral response is a significant challenge. By carefully designing our behavioral protocols we were able to achieve just that. We trained honey bees in such a way that allowed us to show that sensory adaptation reduces appetitive learning of the adapted component of the mixture, while concomitantly enhances learning of the non-adapted minority components that would normally stay occluded if sensory adaptation did not mediate. Thus, our results suggest that sensory adaptation is critical to allow detection of minority components that are present in complex mixtures, emphasizing that it is a fundamental mechanism capable of altering sensitivity to discrete stimuli changing the perceptual quality of a mixture. This phenomenon, despite causing a reduction on the conditioned response as was observed during testing, it caused a significant reduction of the behavioral response during training to the conditioned odorant. The time scale that we observed for the induction of adaptation is in line with previous results (Stortkuhl et al., 1999), although more experiments are necessary to determine the shortest duration capable of eliciting sensory adaptation in honeybees, as well as the minimum recovery period. As we have previously shown, odorant pre-exposure is capable of eliciting both, cis and trans adaptation and importantly, the salience of the odorant has a big role on the adaptation triggered.

We found that in cases in which the concentration of the minority component was such that the animals could still easily detect it (e.g. 2% v/v for the 4:1), adaptation to the majority component was observed as a reduction on the learning of this component, as observed in the testing session. For the two odorants used for these experiments, we found that the ratio 4:1 caused a similar effect, the response to the adapted odorant was reduced compared with the response to the non-adapted odorant for the same animals. On the other hand, the response of the control animals was high for the two components of the mixture, suggesting that at this ratio the animals were able to detect and learn both odorants in the mixture. So, in this case we can conclude that the learning that is taking place during the CS presentation is specific to the minority component that is only present during the mixture presentation and always associated with the sucrose reward. One can think that in this case, the adaptation is functioning like a filter, filtering out non relevant information, i.e. the adapted odorant in the experimental animals.

### Olfactory sensory adaptation improves detection of minority stimuli

We initially expected the animals to have a hard time detecting the minority component, especially for the animals of the control group. Nevertheless, it is clear from these results that at least for the odorants and concentrations used here, the animals are fully capable of detecting both components in the mixture. When the difference in concentration was not big enough (i.e. ratio 4:1) the main effect of sensory adaptation was seen as a reduction on the learning of the adapted odorant. However, the minority component was still in a concentration high enough for the animals to easily detect it, both in the control and adapted groups.

However, when the ratio between the two odorants is such that the minority component is masked by the majority component the observed effect of adaptation is different. In a situation like this, the effect that adaptation is having on the majority component is to reduce its learning, and by doing so, it unmasks the minority component enhancing its detection by the animals. These results highlight the versatility and relevance of this plasticity phenomenon, capable of enhancing the detection power of the animals to allow them to detect relevant environmental stimuli above other stimuli present (as in background segmentation). The adaptation observed in our experiments could originate from a combination of different mechanisms. It could be that the olfactory receptors at the antenna are subject to a very fast and mainly peripheral adaptation. However, it could also be due to another phenomenon, such as habituation (Twick et al., 2014; Wilson and Linster, 2008). Based on results from fruit flies (Stortkuhl et al., 1999) and our behavioral data, these results are consistent with the hypothesis that the phenomenon taking place here is a case of adaptation and not habituation. The time scale used in our experimental protocols also supports this claim. It is important to point out that by means of our behavioral experiments we have shown that this phenomenon reduces the learning of adapted stimuli, while concomitantly enhancing the learning of stimuli that normally would be overshadowed. Additionally, by means of calcium imaging experiments we show that the pattern of activity that encodes a binary mixture is drastically changed following sensory adaptation, in agreement with the behavioral experiments, favoring the encoding or representation of odorants present in lower concentrations. These results show that adaptation is crucial to enhance the sensibility of the animal. It does that by allowing the detection of minor components present in complex mixtures, and by increasing the sensibility of the animal to certain stimuli. In other words, sensory adaptation to an odorant modifies the way the animal perceives a mixture that contains it, favoring the detection of other components of the mixture that might be overshadowed.

## Acknowledgments

F.F.L and N.P. are members of the Argentine National Research Council (CONICET). F.A.G. is funded by a doctoral scholarship by CONICET.

