## Supplemental Material for "Sensory adaptation modulates coding and perceptual quality of odor mixtures"

*contributed equally

#### Supplementary information

To control that the olfactometer reliably delivered a constant concentration of odorant during the 30 seconds pulse we used a PID sensor to measure the delivery of odorant. As Supplementary Figure 1 shows, our system is delivering odorant for the duration of the 30 second pulse without any measurable depletion of the stimulus. An alternative way to analyze the representation of the mixture in relation to the pure components is to calculate the Euclidean distance (ED) between the different activity patterns (Supplementary Table 1). This analysis showed that the distance between the adapted component and the mixture increased after sensory adaptation. However, the ED between the mixture and the non-adapted component, either did not significantly change from its initial value or in one case was significantly reduced. Thus, as we expected, exposure to an odorant moved the representation of the mixture further apart from the exposed component and closer to the non-adapted one.

An interesting observation emerged from comparing the results based on Pearson's correlation and the results based on ED. Both strategies were highly consistent in that after pre-exposure the representation of the mixture moves away from the representation of the exposed odor and closer to the non-exposed odor. However, while the correlation analysis showed that the mixture pattern is almost completely restored after one minute rest period, the ED reveals that such recovery is not complete. These discrepancies might be explained by the intrinsic differences on how correlation and distances are calculated. The correlation analysis between two patterns considers only the relative contribution of each glomerulus to the pattern. In contrast, the ED sums the differences in the absolute activation of each glomerulus across both patterns. These differences suggest that one minute rest period might have been time enough for the mixture to recover its original pattern in regards to the relative contribution of each glomerulus, but not its original response values.

To further evaluate if sensory adaptation affected the response of the non-adapted odorant, we plotted the correlation coefficient between the minor component (i.e., odorant B) and the different mixture presentations as a function of the correlation coefficient between the major component and the different mixture presentations (Supplementary Figure 3). This analysis supports the idea that the sensory adaptation elicited by different odorants is not equal. One could expect that when an odorant is pre-exposed, its correlation with the mixture will change across the axis representing that odorant alone, and will not change on the dimension of the other odorant. However, the effect of adapting to acetophenone caused a global change that affected all the activated glomeruli. On the other hand, when the odorant used to trigger adaptation is 1-hexanol, the change in the acetophenone response is very small. In summary, these results support the idea that sensory adaptation elicited by different stimuli results in different degrees of reduction of the response.

#### Supplementary Figures Legends

**Supplementary Figure 1. Custom made olfactometer reliably delivers odorant for 30 seconds.** Example traces of a PID recording of two 30 second pulses of 8% v/v acetophenone (**A**) and 8% v/v 1-hexanol (**B**). The figure clearly shows that the olfactometer delivers odorant molecules for the duration of the 30 second pulses.

**Supplementary Figure 2. Olfactory adaptation alters the representation of odor mixtures in the antennal lobe.** Spatiotemporal patterns of the two pure odorants and the three mixture presentations across all PNs are displayed in a PC space. **A.** Animals were adapted to acetophenone and the ratio to 1-hexanol was 4:1. **B.** Animals were adapted to 1-hexanol and the ratio to acetophenone was 4:1. **C** and **D.** Same as in A and B but the ratio between odorants was 40:1.

**Supplementary Figure 3. Sensory adaptation depends on the salience of stimuli. A.** Plot showing the Pearson correlation coefficient between the odorant B (minority component) and the different mixtures as a function of the Pearson correlation coefficient between the odorant A (majority component) and the different mixtures. **A.** Animals were adapted to acetophenone and the ratio to 1-hexanol was 4:1. **B.** Animals were adapted to 1-hexanol and the ratio to acetophenone was 4:1. **C** and **D.** Same as in A and B but the ratio between odorants was 40:1. Data is presented as mean ± SEM.

**Supplementary Table 1. Olfactory adaptation causes Euclidean Distances between odorants and the different mixtures to be modulated**.

| | **ACE-MIX PRE** | **ACE-MIX ADAPT** | **ACE-MIX RECOV** | **1-HEX-MIX PRE** | **1-HEX-MIX ADAPT** | **1-HEX-MIX RECOV** | **ACE-1-HEX** |
|---|---|---|---|---|---|---|---|
| **ACE 4:1** | 3.26 ± 0.67$^{C}$ | 12.43 ± 1.81$^{B}$ | 7.84 ± 0.89$^{A}$ | 5.69 ± 0.77$^{A,C}$ | 8.49 ± 1.03$^{A}$ | 5.44 ± 1.27$^{A,C}$ | 6.39 ± 0.67$^{A, C}$ |
| **ACE 40:1** | 3.29 ± 0.62$^{B}$ | 17.79 ± 5.42$^{A}$ | 9.91 ± 3.63$^{A,B}$ | 17.82 ± 3.91$^{A}$ | 10.36 ± 3.76$^{A,B}$ | 9.72 ± 3.00$^{A,B}$ | 16.18 ± 3.23$^{A}$ |
| | **ACE-MIX PRE** | **ACE-MIX ADAPT** | **ACE-MIX POST** | **1-HEX-MIX PRE** | **1-HEX-MIX ADAPT** | **1-HEX-MIX RECOV** | **1-HEX-ACE** |
| **HEX 4:1** | 4.87 ± 1.24$^{A,B}$ | 5.10 ± 1.53$^{A,B}$ | 3.36 ± 0.54$^{A}$ | 2.61 ± 0.38$^{A}$ | 7.74 ± 1.92$^{B}$ | 3.43 ± 0.78$^{A}$ | 4.58 ± 0.82$^{A,B}$ |
| **HEX 40:1** | 19.87 ± 1.47$^{B}$ | 6.17 ± 1.57$^{A}$ | 9.53 ± 3.12$^{A,B,C}$ | 6.05 ± 2.55$^{A}$ | 17.43 ± 2.45$^{B,C}$ | 7.64 ± 1.76$^{A,C}$ | 15.15 ± 2.60$^{A,B,C}$ |
| | **OCT-MIX PRE** | **OCT-MIX ADAPT** | **OCT-MIX RECOV** | **GER-MIX PRE** | **GER-MIX ADAPT** | **GER-MIX RECOV** | **OCT-GER** |
| **OCT 1:1** | 5.81 ± 0.36$^{A}$ | 13.24 ± 2.49$^{B}$ | 5.56 ± 0.99$^{A}$ | 12.81 ± 2.68$^{A,B}$ | 8.84 ± 1.72$^{A,B}$ | 12.01 ± 2.79$^{A,B}$ | 15.34 ± 3.28$^{B}$ |

Data was analyzed by means of Repeated Measured One-way ANOVA, $p<0.01$. Different letters represent significant differences by Tukey's multiple comparisons, $p < 0.01$.

### Supplementary Figure 1

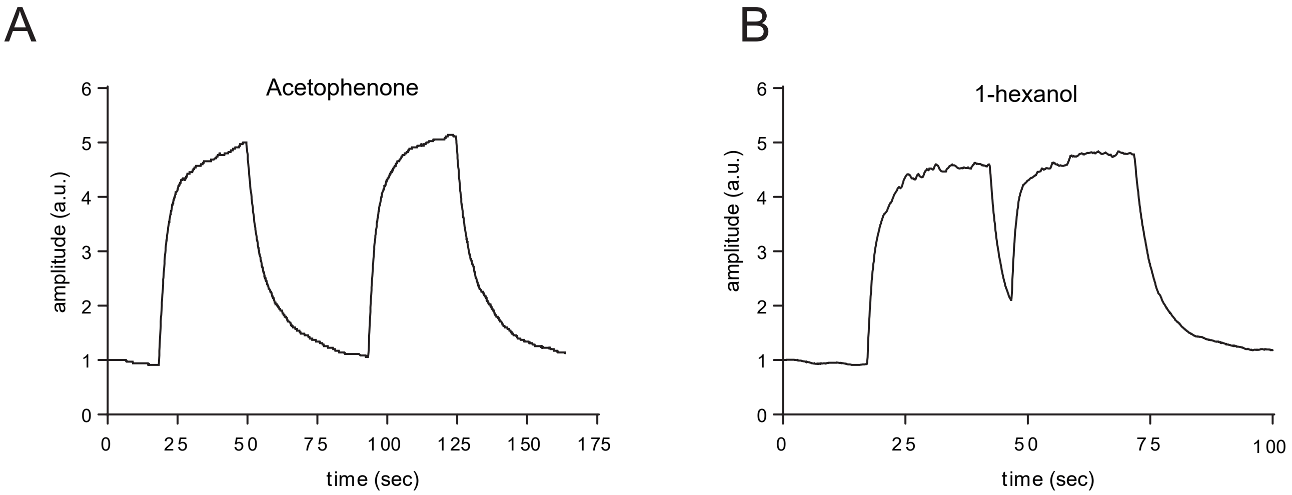

### Supplementary Figure 2

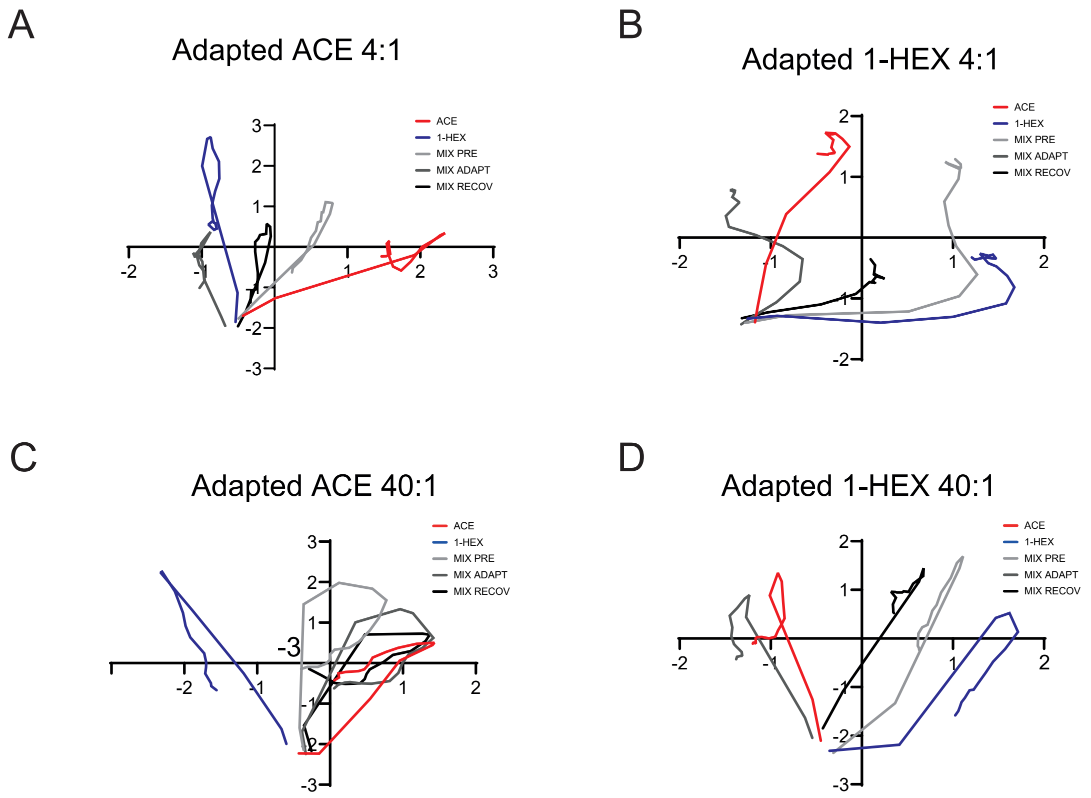

### Supplementary Figure 3

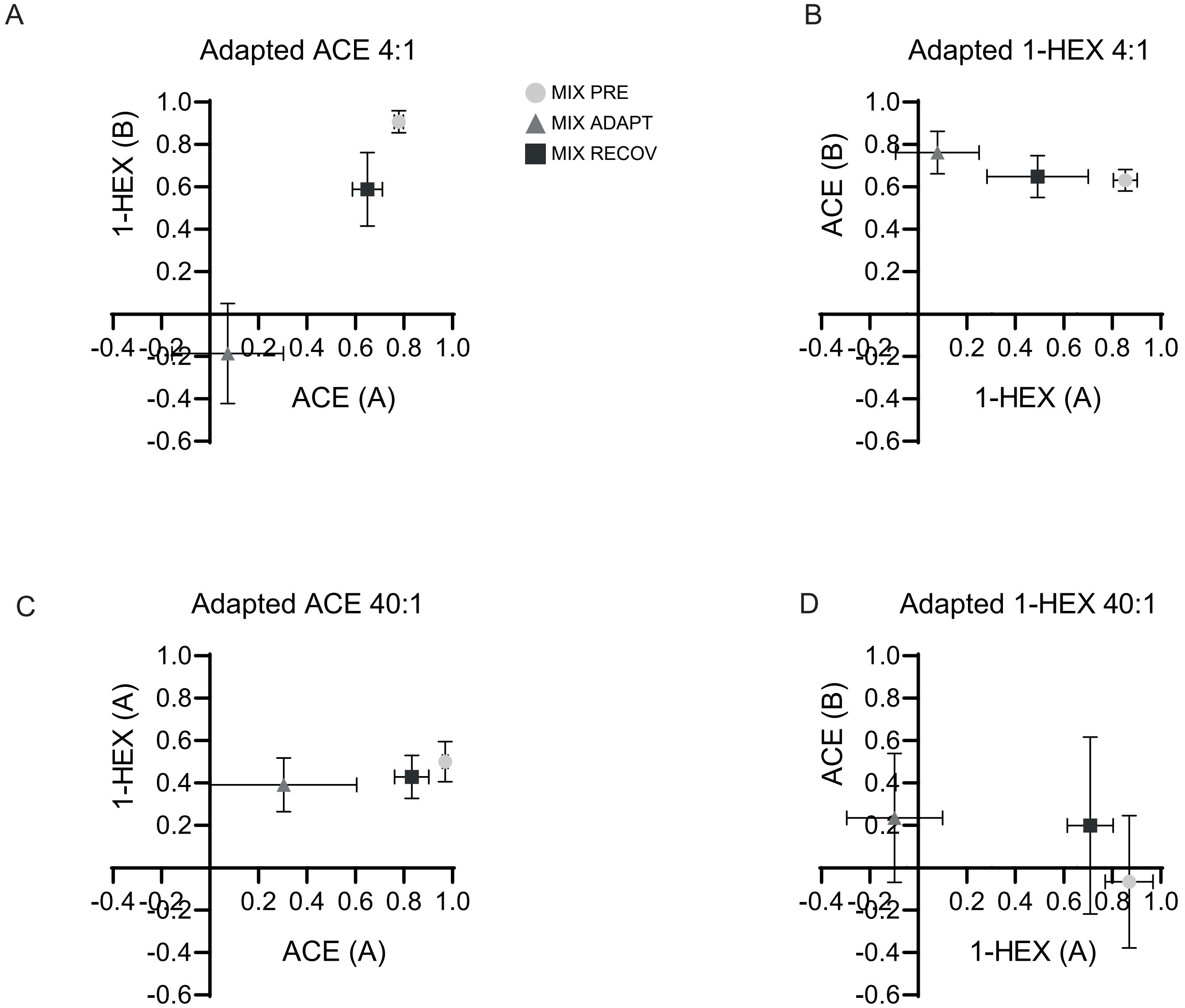
